# Boundary-guided cell alignment drives mouse epiblast maturation

**DOI:** 10.1101/2025.04.28.650859

**Authors:** Takafumi Ichikawa, Pamela C. Guruciaga, Shuchang Hu, Steffen Plunder, Mei Makino, Marina Hamaji, Anniek Stokkermans, Shinjiro Yoshida, Anna Erzberger, Takashi Hiiragi

**Author notes:** Correspondence should be addressed to T.I., A.E. and T.H. These authors contributed equally to this work (Takafumi Ichikawa, Pamela C. Guruciaga). Developmental Biology Unit, European Molecular Biology Laboratory, 69117 Heidelberg, Germany (Shuchang Hu).

## Abstract

Symmetry breaking and pattern formation are critical events that occur throughout embryonic development. In early mouse development, a mass of non-polarized epiblast (EPI) cells in the blastocyst forms the egg-cylinder, while cells become apico-basally polarized and build a radial configuration. Yet, what drives the formation of this tissue architecture remains unclear. Here, we demonstrate that orientational patterning of EPI cells is dictated by heterogeneous tissue boundaries, which then defines central lumen positioning. We show that EPI cells progressively orient perpendicular to the visceral endoderm (VE) boundary enriched with laminin and active integrin β1, but parallel to the extraembryonic ectoderm interface. These orientation dynamics are consistent with general boundary-induced alignment effects in polar materials, with a topological defect predicting the position where the pro-amniotic cavity nucleates. Knockout of laminin γ1 and integrin β1 confirms the essential role of adhesion at the EPI-VE-boundary. The established EPI pattern, in turn, facilitates ERK activation to ensure proper EPI maturation. Together, these findings present the mechanistic basis and functional significance of EPI tissue patterning.

## Introduction

During embryonic development and homeostasis, cells and tissues repeatedly break symmetry and form patterns. At the tissue scale, patterns can emerge through biochemical and mechanical interactions between cells or at the supracellular level. Morphogen signaling, for example, instructs the formation of tissue patterns with distinct material properties (Briscoe and Small 2015; Pinheiro et al. 2022; Wolpert 1969; Yang et al. 2023). While these mechanisms have been studied, the effect of tissue boundaries on tissue patterning remains less understood (Huycke et al. 2024; Landsberg et al. 2009; Martyn and Gartner 2021; Monier et al. 2010), particularly when properties of these boundaries are heterogeneous. Recent studies have begun to investigate how interactions between cells and the extracellular matrix (ECM) deposited at tissue boundaries contribute to tissue formation (Munjal et al. 2021; Palmquist et al. 2022). However, specific mechanisms by which cell-ECM interactions influence cell arrangement, tissue patterning, and their subsequent functional consequences remain elusive.

Orientational order —a spatial pattern in living systems— emerges at various scales, from subcellular structures to whole organisms (Doostmohammadi and Ladoux 2022; Maroudas-Sacks et al. 2021; Sanchez et al. 2012). Directional cellular organization, analogous to nematic ordering in liquid crystals (de Gennes and Prost 1995), has been observed in two-dimensional (2D) cell cultures, such as neural progenitors (Kawaguchi, Kageyama, and Sano 2017) and epithelial cells (Saw et al. 2017). However, our understanding of three-dimensional (3D) cell arrangement in tissues remains limited, primarily due to the technical challenges associated with monitoring cellular dynamics within a 3D tissue undergoing pattern formation. Moreover, investigating the potential impact of cell-ECM interactions at the tissue boundary requires approaches to manipulate ECM deposition without disrupting overall tissue architecture.

The early mouse embryo provides a model system to investigate the mechanisms of tissue pattern formation (Bedzhov and Zernicka-Goetz 2014; Moghe et al. 2025; Saiz et al. 2016). Mammalian embryos are derived from the epiblast (EPI) that forms in the blastocyst as an aggregate of non-polarized cells (Chazaud et al. 2006; Moghe et al. 2025; Ohnishi et al. 2014; Plusa et al. 2008; Yanagida et al. 2022). Upon implantation and by embryonic day 5.5 (E5.5) — at the cellular level— EPI cells undergo elongation, acquiring apico-basal polarity, and arrange in a radial manner, while —at the tissue level— the EPI tissue transforms into the cup-shaped egg-cylinder structure (Bedzhov and Zernicka-Goetz 2014; Ichikawa et al. 2022; Y. S. Kim et al. 2021; Molè et al. 2021). During this morphogenesis, the EPI is enveloped by two extraembryonic tissues: the primitive endoderm (PrE), which differentiates into the visceral endoderm (VE), and the polar trophectoderm (pTE), which gives rise to the extraembryonic ectoderm (ExE). These interactions establish two distinct tissue boundaries with potentially different properties. Indeed, the precise cellular and molecular mechanisms underlying EPI patterning remain elusive, largely due to limited access to the dynamic cellular processes occurring within the uterine tissue.

In this study, building upon our recently established method for *ex vivo* embryo culture and live-imaging (Ichikawa et al. 2022), together with quantitative image analyses and a theory for boundary-driven polar ordering, we investigate how the EPI tissue pattern is established during mouse peri-implantation development.

## Results

### EPI cells progressively align their long axes with each other and against the tissue boundaries

To understand cellular mechanisms underlying EPI tissue patterning, we first quantitatively characterized EPI cell orientation in mouse embryos developed *in utero*, at stages from E4.5 to E5.25. To this end, we developed an image analysis pipeline that semi-automatically segments EPI cell membranes based on the combined signal of E-cadherin and phalloidin immunofluorescence (Figures S1A and S1B). This analysis identifies the cellular long axis in three dimensions (3D) (Figure 1A). Using this data, we examined the relationship between the long axes of neighboring EPI cells as well as between EPI cells and the EPI tissue boundaries in 3D. Measurement of the angle between the long axes of neighboring cells showed that the angle decreases as the embryo develops with an increasing number of EPI cells. This observation indicates that elongated EPI cells progressively align their long axes with each other during egg-cylinder formation (Figure 1B), suggesting that cell-cell interactions may drive this alignment (Ichikawa et al. 2022). In addition, EPI cells establish specific orientations relative to the two distinct tissue-tissue boundaries, namely with the polar trophectoderm (pTE) or extraembryonic ectoderm (ExE) (hereafter referred to as “ExE-boundary”), and with the primitive endoderm (PrE) or visceral endoderm (VE) (“VE-boundary”). We measured the angle between the cellular long axis and the normal vector to each tissue boundary (Figures S1C and S1D), while the EPI cell number increased from 15 to 104 between E4.25 and E5.25. The median angle at the ExE-boundary increases from 65.57° to 77.80° (Figure 1C), whereas that at the VE-boundary decreases from 61.53° to 24.50° (Figure 1D). These data show that EPI cells progressively align themselves parallel to the ExE-boundary and perpendicular to the VE-boundary.

**Figure 1.**
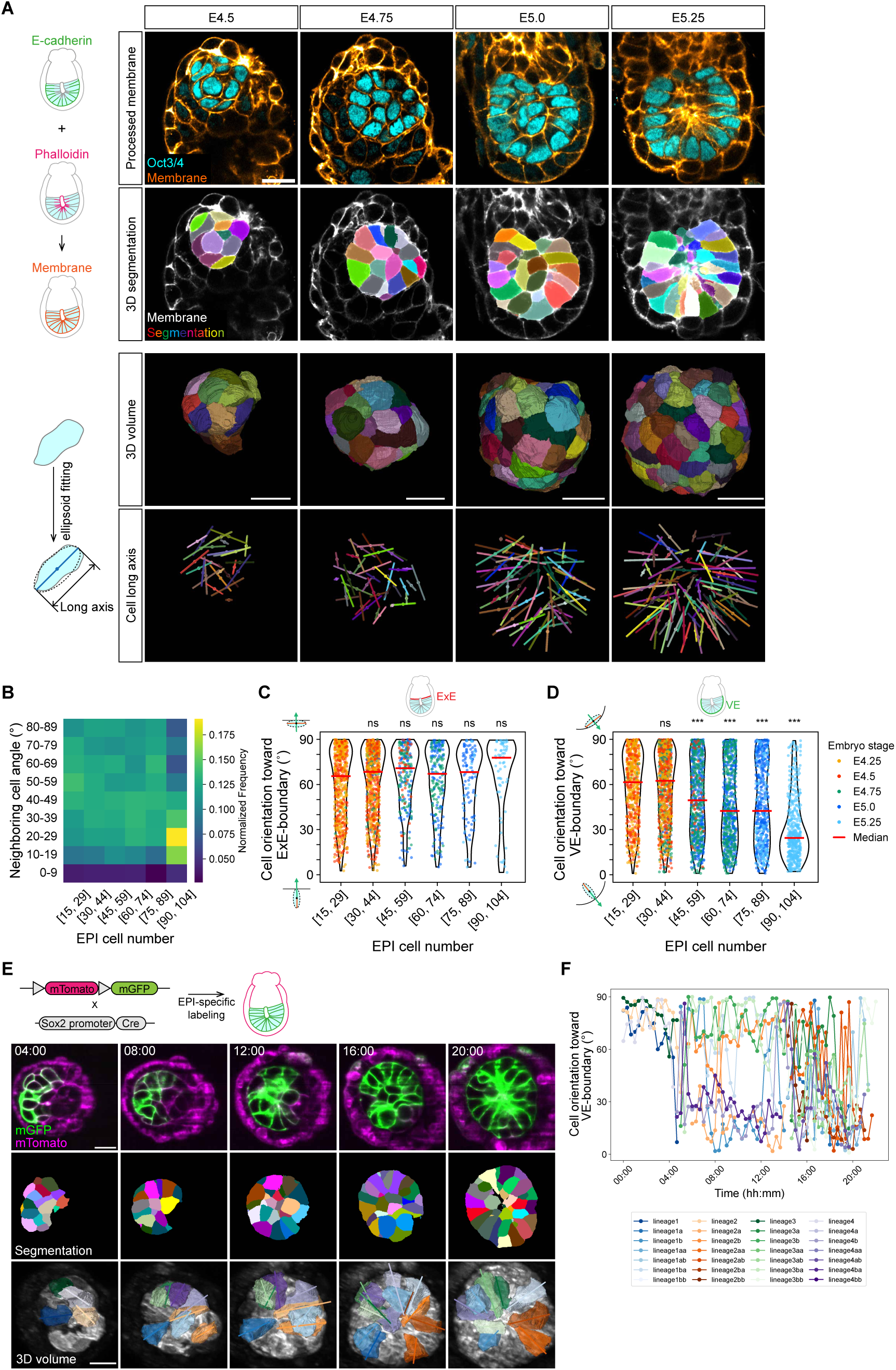
Progressive cell alignment and orientation to the boundary underlie EPI patterning. (A) Immunofluorescence images (top row) and corresponding membrane segmentation (second row) of representative embryos from E4.5 to E5.25, stained for Oct3/4 (EPI) and cell membrane, generated by combining E-cadherin and phalloidin signals. Based on the 3D individual cell volume (third row), cell long axes (bottom row) were extracted by ellipsoid fitting. *n* = 37 (E4.5), 17 (E4.75), 21 (E5.0), and 7 (E5.25) embryos segmented and analyzed from at least three independent embryo recovery experiments. (B) Distribution of angles between the long axes of neighboring cells, represented as smoothed histograms, color-coded based on every 25^th^ percentile. Data from E4.5 to E5.25 embryos are grouped by EPI cell number in intervals of 15. *n* = 36 embryos analyzed for EPI cell number 15-29, 36 for 30-44, 18 for 45-59, 17 for 60-74, 8 for 75-89, and 4 for 90-104. (C-D) Angle measurement between the cell long axis and the normal vector to the tissue boundary, represented as violin plots with individual data points overlaid. Each plot shows the distribution of angles for EPI cells in contact with the ExE-boundary (C) and VE-boundary (D). Dot colors indicate the embryo stage at the collection. Data are grouped by EPI cell number, with group medians shown by red bars. *n* = 36 embryos analyzed for EPI cell number 15-29, 35 for 30-44, 15 for 45-59, 12 for 60-74, 8 for 75-89, and 5 for 90-104. (E) Time-lapse images of representative Sox2-Cre;mTmG embryos developed in 3D-geec. Green, mG (EPI); magenta, mT (other tissues). Time is shown as hh:mm, with t=00:00 marking the start of imaging. *n* = 5 embryos. (F) Angle measurement between the cell long axis and the normal vector to the VE-boundary of tracked cells. Each line represents the measurement from a tracked EPI cell, with line colors indicating cell lineages. Scale bars, 20 µm. See also Figure S1 and Video S1.

These findings were further corroborated by live-imaging of Sox2-Cre;mTmG embryos developing *ex vivo* using the <u>3D</u>-<u>g</u>el <u>e</u>mbedded <u>e</u>mbryo <u>c</u>ulture (3D-geec) (Figure 1E) (Ichikawa et al. 2022). The analysis of cell orientation dynamics demonstrated that EPI cells progressively establish their orientation perpendicular to the VE-boundary (Figure 1F). Taken together, our static and dynamic analyses consistently demonstrate that the progressive alignment of elongated EPI cells with each other and relative to the EPI tissue boundaries establishes the EPI tissue-scale alignment pattern.

### Emergence of tissue-scale orientational order in the EPI near the tissue boundary

To quantify the tissue-scale orientational order, we developed a computational tool to systematically measure cell alignment patterns across the entire EPI tissue (Figure 2A). Leveraging the approximate axial symmetry of the embryo shape along the proximal-distal axis, we analyzed the spatial orientation of EPI cells by examining 36 sections at 10-degree rotational intervals around this axis. Within each section, we characterized cell elongation via directions along the major axis of the cell and anisotropy ranging from zero (for spheres) to nearly one (for highly elongated cells) (see Methods). We then computed the weighted nematic order parameter field by averaging cell orientations weighted by their elongation across all rotational planes (see Methods).

**Figure 2.**
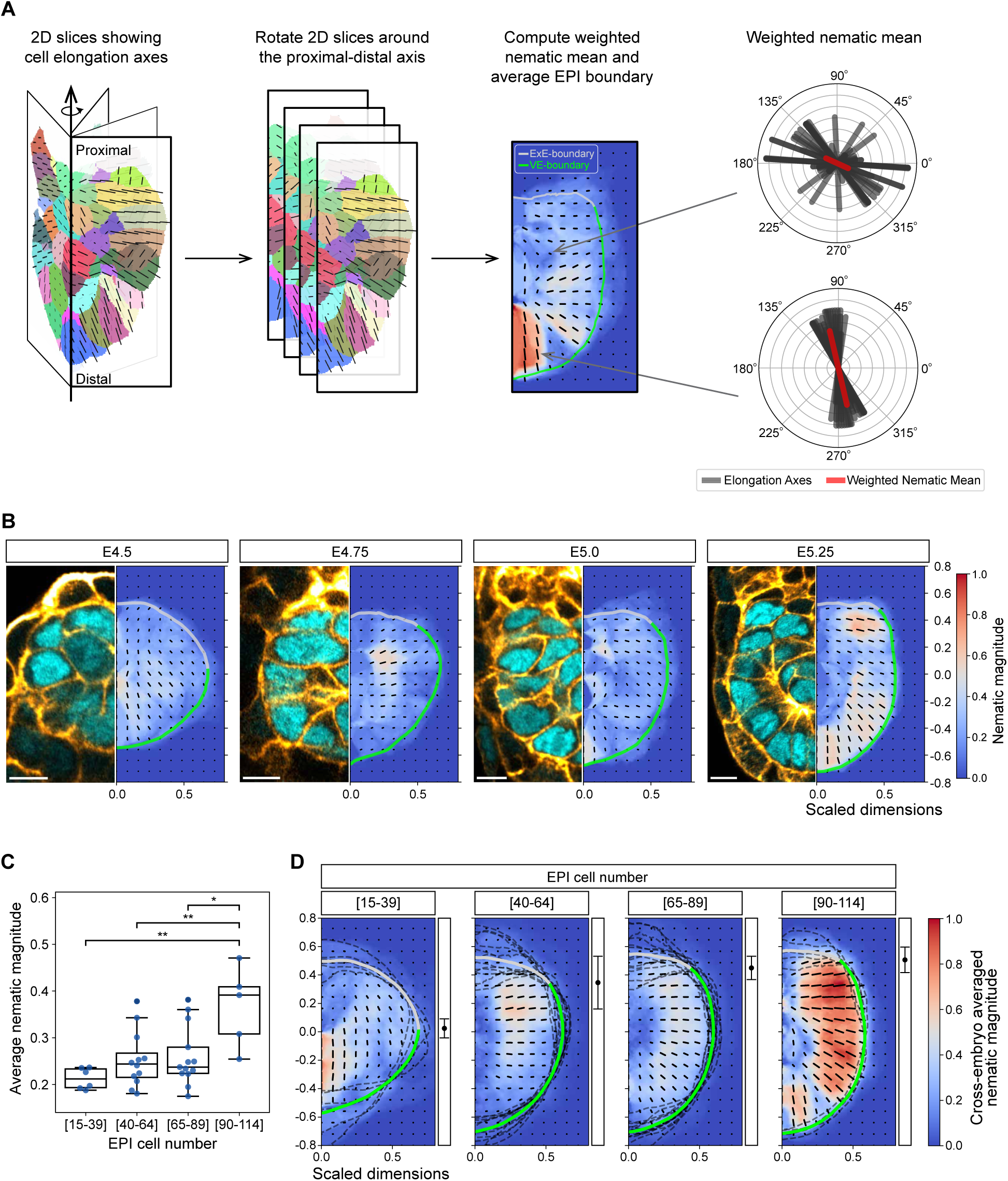
Emergence of tissue-scale cell alignment in the EPI near the tissue boundary. (A) Schematic illustrating the method used to compute tissue-scale cell alignment pattern. An arrow represents the distal-to-proximal axis of the EPI tissue, used as the axis of rotation. Each individual color represents cell segmentation in Figure 1, and the lines indicate orientations along the long axis and lengths reflecting the anisotropy of the cell (left). 36 sections are reconstructed every 10° of rotation around the axis (second from left). With an average EPI boundary shape, the nematic order parameter field was calculated by averaging cell orientations weighted by their elongation across rotational planes (right). Finally, the nematic director field and the magnitude of the nematic alignment were obtained (second from right). (B) Representative embryo images from E4.5 to E5.25 and their corresponding cell alignment map. The color indicates the magnitude of nematic cell alignment, and the lines represent the nematic director vector. Light grey and green thick lines highlight the average ExE and VE-boundaries, respectively. *n* = 6 (E4.5), 9 (E4.75), 16 (E5.0), and 5 (E5.25) embryos analyzed. (C) Box plot of average nematic magnitude within the EPI region, grouped by EPI cell number in embryos from E4.5 to E5.25. Each dot represents an individual embryo. *n*=6 embryos for EPI cell number 15-39, 12 for 40-64, 13 for 65-89, and 5 for 90-114. Tukey’s post-hoc test, \**P*<0.05, \*\**P*<0.01. (D) Cross-embryo average of nematic cell alignment maps grouped by EPI cell number in intervals of 25. The color indicates the magnitude of nematic cell alignment, and the lines represent the mean orientation. Light grey and green thick lines highlight the average ExE and VE-boundaries, respectively, and dark grey broken lines indicate the tissue boundary of each embryo. Black dots with error bars at the right indicate the mean ± SD position of the interface between ExE and VE boundaries along the proximal-distal axis. *n*=6 embryos for EPI cell number 15-39, 12 for 40-64, 13 for 65-89, and 5 for 90-114, used for averaging in each group. Scale bars, 10 µm.

This analysis revealed the emergence of a tissue-scale orientational order within the EPI during development (Figure 2B). In particular, the tissue-scale pattern shows an average perpendicular orientation to the VE-boundary and parallel to the ExE-boundary after E4.75, consistent with our measurements of individual cell orientations (Figures 1C and 1D). We found quantitatively a gradual increase in average nematic magnitude, which measures the degree of cell alignment across the entire EPI region within each embryo (Figure 2C), in agreement with our initial observations (Figure 1). To investigate the progression of alignment patterns while accounting for embryo-to-embryo variability, we averaged the nematic field of EPI tissues grouped by their cell number in intervals of 25 (Figure 2D). Notably, this analysis revealed that the magnitude of nematic alignment is consistently higher near the tissue boundaries compared to the tissue center, despite the variability between embryos. This robust spatial heterogeneity in alignment strength suggests that tissue-tissue boundaries provide critical instructive cues that not only guide local cell orientation but also influence the global organization of the EPI tissue. These findings indicate a boundary-mediated mechanism for establishing EPI cell organization.

### Surface-induced ordering determines the orientational pattern of the EPI tissue

The progressive alignment of elongated cells within the tissue and against the tissue boundaries is reminiscent of how anisotropic particles develop orientational order through surface interactions (Crawford, Stannarius, and Doane 1991; Jerome 1991; Sheng 1976, 1982). While EPI cells establish the radial orientation, EPI cells become polarized with basal domains attaching to the ECM (Bedzhov and Zernicka-Goetz 2014; Ichikawa et al. 2022; Molè et al. 2021). These similarities between polar cell alignment in developing tissues and surface-induced ordering of polar materials prompted us to investigate whether the theoretical framework of surface-induced ordering could provide insights into the mechanism of EPI tissue patterning.

Using the Landau-de Gennes approach, which describes orientational order in anisotropic materials (de Gennes and Prost 1995), we conceptualized EPI cells as polar particles in a 3D space, constituting a polar fluid characterized by a vector order parameter field p. This local order parameter takes a value according to the strength of aligning interactions within the bulk, which competes with the tendency to align with specific directions promoted by the boundaries (Guruciaga et al. 2024; Prinsen and van der Schoot 2003; Seyednejad, Mozaffari, and Ejtehadi 2013). Specifically, we consider an effective free energy functional that takes into account the tendency of cells to align with their neighbors (Pallarès et al. 2022; Pérez-González et al. 2019), modulated by the correlation length *ξ*. In addition, it incorporates phenomenologically the interactions between EPI cells and the boundaries via a soft surface anchoring (Ravnik and Žumer 2009; Seyednejad, Mozaffari, and Ejtehadi 2013), which penalizes deviations from a preferred value with a strength given by the anchoring length *λ* (see Methods). Based on our experimental observations, we implemented perpendicular and parallel anchoring terms for the VE- and ExE-boundaries, respectively (Figure 3A). To examine how alignment arises within a geometry that corresponds to that of the EPI tissue, we constructed axially symmetric shapes formed by two spherical shells representing ExE- and VE-boundaries (Figure 3B, see Methods). By minimizing the free energy functional for different correlation and anchoring values, we obtained the corresponding order parameter fields, revealing a critical transition in the alignment pattern when surface anchoring overcomes the cost of bulk distortions. In this surface-dominated regime, singular points (topological defects) appear in the order parameter field, and the global degree of order increases significantly (Figure 3C) (Guruciaga et al. 2024).

**Figure 3.**
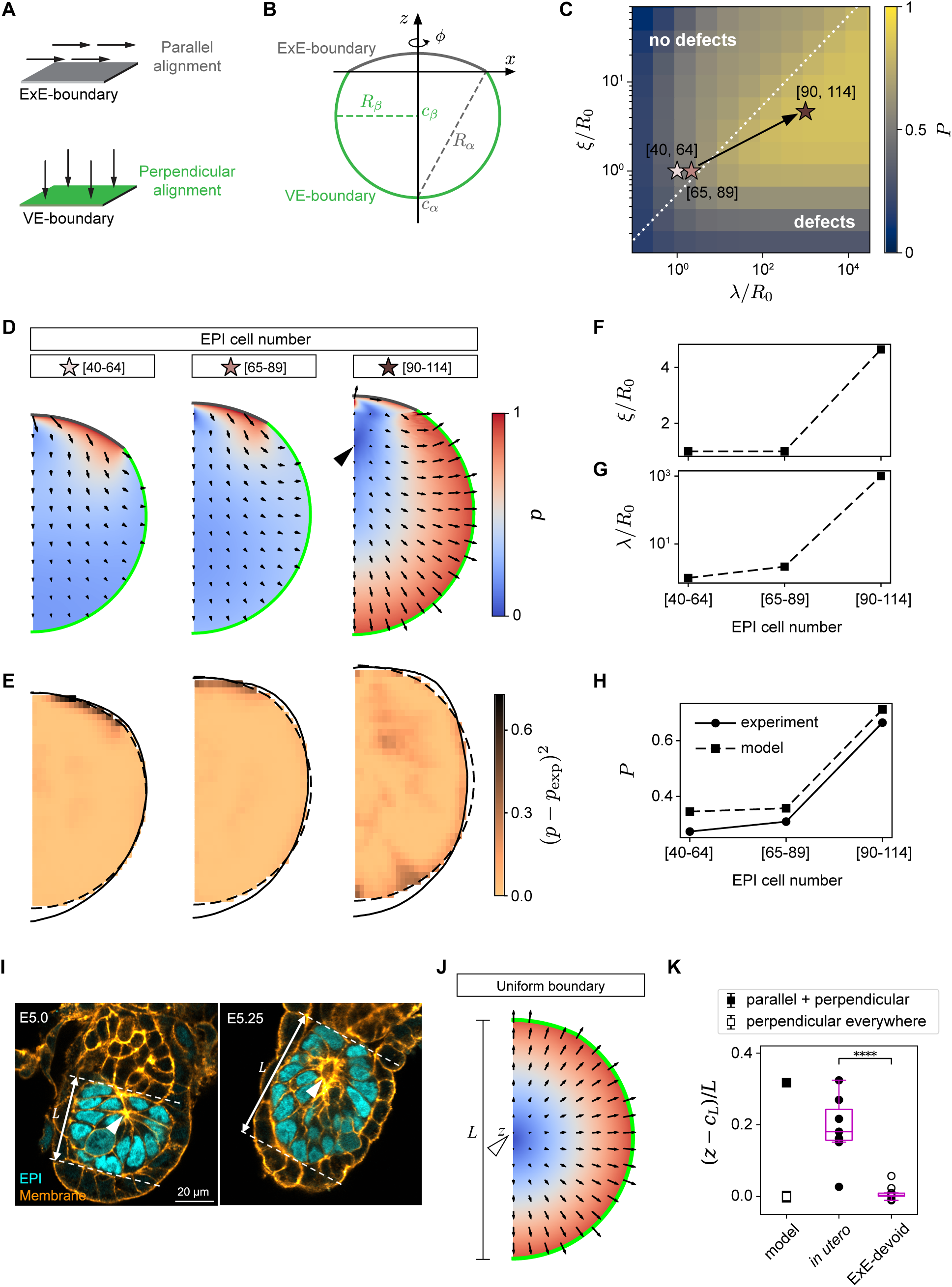
Progressive EPI ordering is driven by boundary-induced alignment. (A) Schematic illustrating two different surfaces that favor distinct cell alignments: tangential alignment at the ExE-boundary (grey) and perpendicular alignment at the VE-boundary (green). (B) Schematic representation of *in silico* geometry based on the EPI tissue. Axially symmetric EPI tissue shape was represented by two spherical shells *S*_*μ*_, *μ* = *⍺*, *β* with radius *R*_*µ*_ and center *c*_*µ*_. These geometrical parameters are obtained by fitting the ExE-boundary (dark grey) and the VE-boundary (dark green) data obtained in Figure 2D. (C) We computed the degree of global order *P* of the order parameter field minimizing the effective free energy as a function of the correlation and anchoring lengths (*ξ* and λ) relative to the characteristic system size *R*_0_ = (3*V*_0_/4*π*)^1/3^, where *V*_0_ is the EPI tissue volume. Stars show the position in the parameter space of the three developmental timepoints studied obtained by parameter fitting (D-G). The dotted line marks the transition between defect-free and defect-containing regimes. (D) Order parameter fields **p** for the best set of parameters for each embryo stage based on EPI cell numbers. The color map shows their magnitude *p* = |**p**|. Black arrowhead marks the position of the topological defect. (E) Square difference between the magnitude of the experimentally determined elongation field, *p*_exp_, and that of the fitted order parameter field, *p*, at each point. (F-G) Material parameters obtained by fitting the theoretical field to the experimentally determined cell orientation field, plotted against the stage of embryos grouped by EPI cell numbers. Both the correlation length *ξ* (F) and the anchoring length *λ* (G) relative to the characeristic system size *R*_0_ increase with the number of cells. (H) The degree of global order *P* grows with the EPI stage both in the model and in the experiments. (I) Immunofluorescence images of representative embryos, stained for Oct3/4 (EPI; cyan), and cell membrane (orange). White arrowheads indicate rosette structures or a nascent lumen. Tissue length *L* is measured as the distance between the EPI-ExE interface and the distal tip. (J) Order parameter field calculated with the geometry and material parameters of the embryo stage at which the EPI cell number ranges from 90 to 114, with a uniform perpendicular boundary preference. White arrowhead marks the position of the topological defect (at position z along the system height L). (K) Quantification of a topological defect (model) and a lumen (*in utero* and ExE-devoid embryos (data adapted from (Guruciaga et al. 2024))) position along the proximal-distal axis, relative to the tissue center (*c*_*L*_ = *L*/2) and scaled by tissue length *L*. Scale bars, 20 µm.

To quantify the relative strengths of bulk and surface interactions in the EPI tissue, we fitted the correlation and anchoring length parameters using the measured cell elongation pattern (Figure 2D) as an estimate of the order parameter magnitude at each position, but withholding the measured average orientations for a subsequent parameter-free comparison. Specifically, we defined the experimental polarity magnitude *p*_exp_ as the strength of the nematic alignment normalized by its maximum across developmental stages and identified the parameter values that minimized the difference between *p*_exp_ and the theoretical magnitude *p* = |p| for each stage (Figures 3D and 3E, see Methods). The theoretical results were overall in good agreement with the experimental measurements, with a close correspondence in the time-course of the global degree of order across different embryonic stages. Notably, by the time the EPI cell number reached 90 to 114 cells, the local direction of the fitted order parameter field was close to that of the corresponding experimental orientation field (Figures 2D and 3E), despite having been fitted using only the magnitude.

### Maturation of boundary anchoring triggers lumen nucleation

We found that both the correlation and anchoring length estimates increased over the course of embryo development, with the increase in the latter much stronger than in the former (Figures 3F and 3G). These results suggest that while both cell-cell interactions and cell-boundary interactions strengthen over time, the boundary interactions become dominant at the later stage. Correspondingly, the degree of global order increased from early to late stages in both theory and experiments (Figure 3H). Moreover, our parameter estimates indicate that the EPI crosses the transition into the surface-dominated regime (Figure 3C). Given that topological defect positions mark the locations of apical domain clustering and lumen nucleation (Guruciaga et al. 2024), these results suggest that rosettes and lumen initiation sites within the EPI should appear once cell-boundary interactions become sufficiently strong to overcome cell-cell alignment effects, i.e. that the maturation of cell-boundary interactions triggers lumen initiation. More specifically, we predict that molecular determinants of boundary anchoring that mediate cell-ECM interactions should correspondingly increase over the course of development, achieving lumen nucleation at the appropriate developmental stage. Supporting this notion, we consistently observed lumen nucleation once the EPI cell number reached approximately 100 cells (Figure 3I) (Ichikawa et al. 2022).

### Boundary heterogeneity guides lumen positioning

Using these experimental estimations based on material parameters, we next investigated how boundary *heterogeneity* affects internal organization. Specifically, we compared order parameter fields with and without an ExE-like interface, the latter having uniform perpendicular anchoring at entire boundaries while maintaining other parameters the same. Our simulation reveals that the position of topological defects consistently shifted towards the ExE-boundary in the presence of non-uniform anchoring (Figures 3D and 3J), predicting a similar shift in rosette formations and subsequent lumen initiation sites in the EPI tissue. Our experimental observations confirmed this prediction, showing rosette and lumen positions were indeed shifted towards the ExE-boundary (Figure 3K), as opposed to the near-center lumina observed in the EPI tissue entirely surrounded by the VE, where the ExE had been removed by immunosurgery (Guruciaga et al. 2024). Hence, these findings demonstrate that spatial heterogeneity in boundary properties plays a crucial role in guiding tissue architecture in developing embryos.

### Integrin-ECM adhesion is specifically established at the VE-boundary

Having established that boundary properties and their heterogeneity influence EPI organization, we next sought to identify the molecular mechanisms that create distinct boundary characteristics. Specifically, the perpendicular alignment of the elongated EPI cells at the VE-boundary suggests the presence of ECM components that mediate integrin-based adhesion at the basal domain, such as Collagen IV (Bedzhov and Zernicka-Goetz 2014; Ichikawa et al. 2022; Kyprianou et al. 2020). To comprehensively characterize the molecular difference between the ExE- and VE-boundaries, we first analyzed our single-cell transcriptome data (Bondarenko et al. 2023). Genes expressed at least two-fold higher in PrE or VE cells in comparison to TE or ExE cells included *Lama1*, *Lamb1*, and *Lamc1*, as well as *Col4a1* and *Col4a2* (Figure S2A).

Next, at the protein level, Col4a2-eGFP (Wuergezhen et al. 2025) embryos developing in 3D-geec showed GFP signal accumulation at the VE-boundary while it was lost at the ExE-boundary (Figure S2B). Immunostaining of pan-laminin also showed its progressive enrichment at the VE-boundary and diminishing signal at the ExE-boundary (Figures 4A and 4B). Laminin chain-specific antibodies further confirmed that laminin α1, laminin β1, and laminin γ1 accumulated at the VE-boundary, whereas laminin α5 was abundant within the EPI tissue (Figure S2C), in line with our transcriptome analysis (Figure S2A). Moreover, the intensity of the local laminin signal was inversely correlated with the angle between the EPI cell long-axis and the VE-boundary normal (Figure 4C), supporting a key role for ECM in guiding EPI cell orientation. Consistently, an active form of a major laminin receptor subunit integrin β1 became enriched at the VE-boundary (Figures 4D and 4E), again in correlation with EPI cell orientation (Figure 4F). Collectively, these data demonstrate that integrin-ECM adhesion is specifically established at the VE-boundary, where EPI cells adopt their perpendicular orientation. This suggests that integrin-ECM interactions serve to anchor cells at the boundary and guide cellular alignment, thereby contributing to tissue-scale organization.

**Figure 4.**
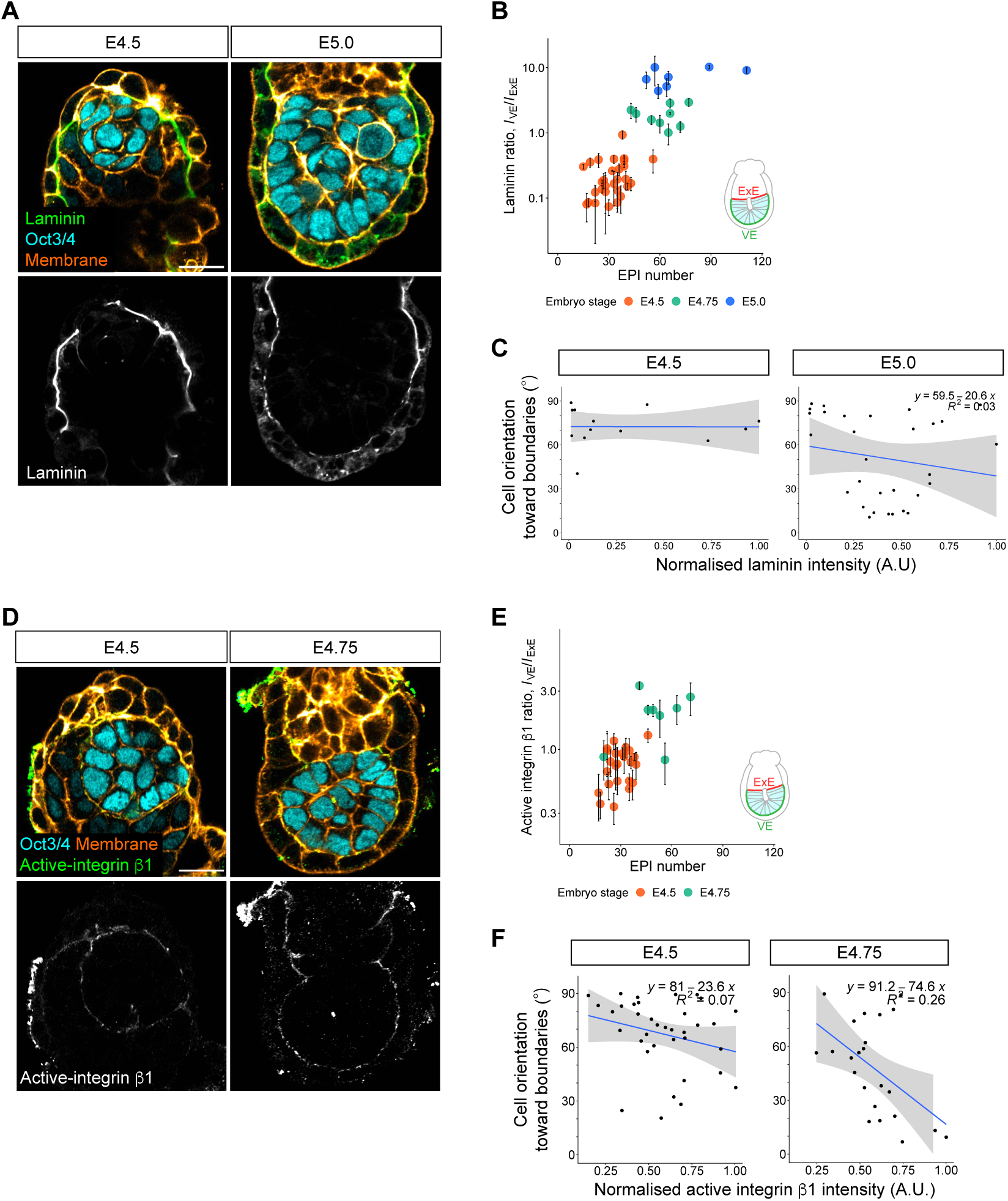
Integrin-laminin adhesion is specifically established at the VE-boundary, correlating with the EPI cell orientation. (A) Immunofluorescence images of representative embryos from E4.5 to E5.0, stained for laminin (green), Oct3/4 (EPI; cyan), and cell membrane (orange). *n* = 29 (E4.5), 9 (E4.75), and 7 (E5.0) embryos analyzed from at least three independent embryo recovery experiments. (B) Quantification of laminin distribution at the tissue boundary, shown as the ratio of intensity at the VE-boundary to that at the ExE-boundary, based on images in (A). Data are presented as mean ± SD. Each dot represents an individual embryo, plotted against EPI cell number. (C) Scatter plot of EPI cell orientation with respect to surface normal versus laminin intensity at the local tissue boundary, based on images in (A). Cells on the middle plane in the 3D EPI volume were used for the analysis. Each dot represents an individual EPI cell in contact with the tissue boundary. Blue line, linear regression. (D) Immunofluorescence images of representative embryos from E4.5 to E4.75, stained for active integrin β1 (green), Oct3/4 (EPI; cyan), and cell membrane. *n* = 24 (E4.5) and 8 (E4.75) embryos analyzed from at least three independent embryo recovery experiments. (E) Quantification of active integrin β1 distribution at the tissue boundary, shown as the ratio of intensity at the VE-boundary to that at the ExE-boundary, based on images in (D). Data are presented as mean ± SD. Each dot represents an individual embryo, plotted against EPI cell number. (F) Scatter plot of EPI cell orientation versus active integrin β1 intensity at the local tissue boundary, based on images in (D). Cells on the middle plane in the 3D EPI volume were used for the analysis. Each dot represents an individual EPI cell in contact with the tissue boundary. Blue line, linear regression. Scale bars, 20 µm. See also Figure S2.

### EPI ordering is dependent on laminin γ1 and integrin β1 anchoring

To further investigate the functional role of boundaries in EPI ordering, we first tested *in silico* the impact of reducing the anchoring strength at the boundary without changing tissue shape (Figure 5A). Loss of anchoring resulted in the disruption of the polarity alignment, i.e. the loss of the orientational order, suggesting the essential role of anchoring to boundaries.

**Figure 5.**
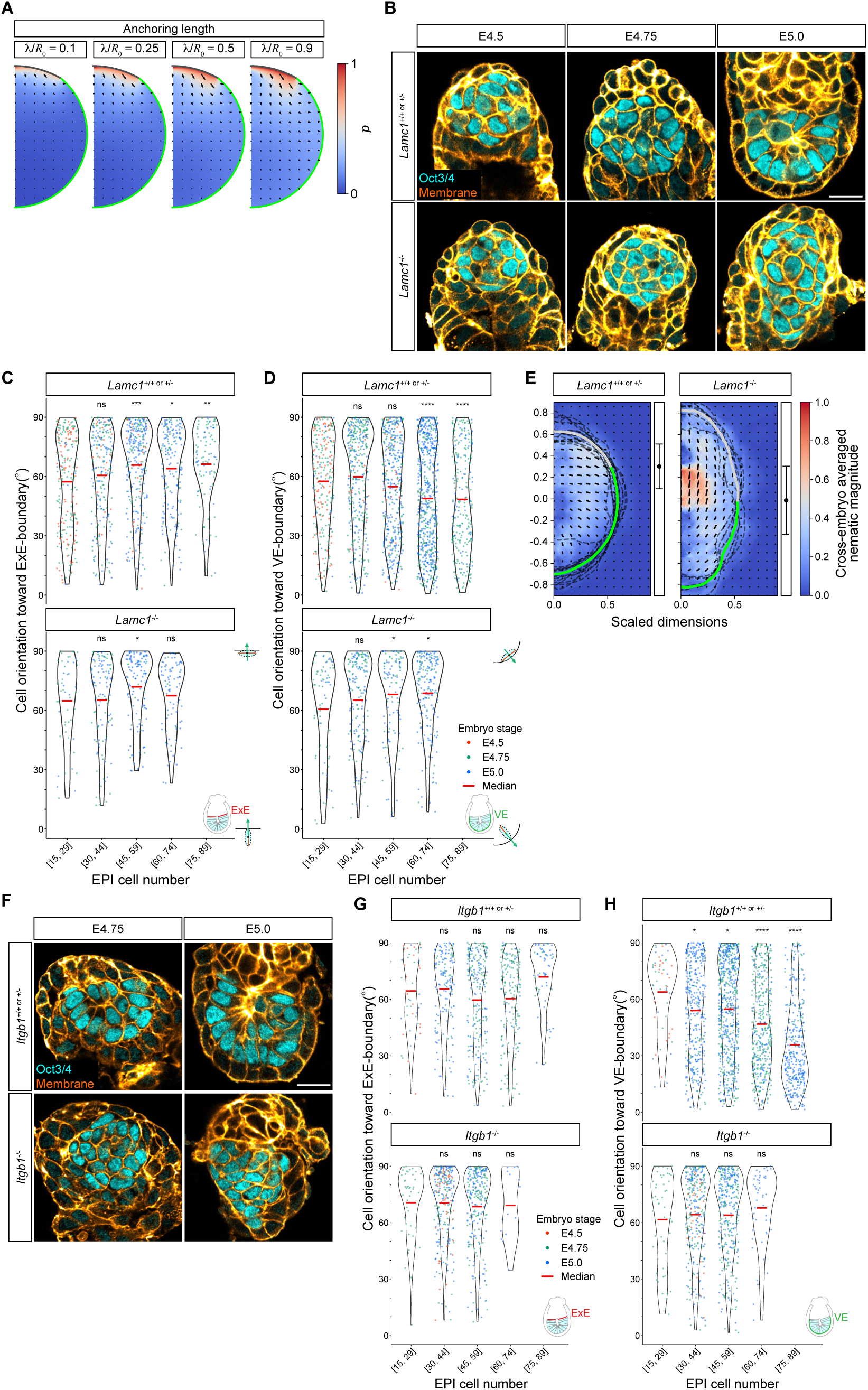
Laminin γ1 and integrin β1 are essential for EPI to build a tissue pattern. (A) Order parameter fields with weak anchoring to the surface characterised by lengths *λ*/*R*_0_ < 1.0 for the average boundary geometry of EPI cell number stage, 65-89. (B) Immunofluorescence images of representative *Lamc1*^+/+ or +/−^ and *Lamc1*^−/−^ embryos from E4.5 to E5.0, stained for Oct3/4 (EPI, cyan) and cell membrane (orange), derived from E-cadherin and phalloidin. *n* = 23 (E4.5), 38 (E4.75), and 65 (E5.0) for *Lamc1*^+/+ or +/−^ and 8 (E4.5), 10 (E4.75), and 20 (E5.0) for *Lamc1^−/−^* embryos examined from at least four independent embryo recovery experiments. (C-D) Angle measurement between the cell long axis and the normal vector to the tissue boundary, represented as violin plots with individual data points overlaid. Each plot shows the distribution of angles for EPI cells in contact with the ExE-boundary (C) and VE-boundary (D). Dot colors indicate the embryo stage at the collection. Data are grouped by EPI cell number, with group means shown by red bars. *n* = 12 (E4.5), 16 (E4.75), and 26 (E5.0) for *Lamc1*^+/+ or +/−^ and 2 (E4.5), 6 (E4.75), and 11 (E5.0) for *Lamc1^−/−^* embryos analyzed from at least four independent embryo recovery experiments. (E) Cross-embryo average of nematic cell alignment maps for *Lamc1*^+/+ or +/−^ and *Lamc1*^−/−^ embryos. The color indicates the magnitude of nematic cell alignment, and the lines represent the mean orientation. Light grey and green thick lines highlight the average ExE and VE-boundaries, respectively, and dark grey broken lines indicate the tissue boundary of each embryo. Black dots with error bars at the right indicate the mean ± SD position of the interface between ExE and VE boundaries along the proximal-distal axis. *n*=15 embryos for *Lamc1*^+/+ or +/−^ and 6 embryos for *Lamc1*^−/−^ mutants. (F) Immunofluorescence images of representative *Itgb1*^+/+ or +/−^ and *Itgb1*^−/−^ embryos from E4.75 to E5.0, stained for Oct3/4 (EPI, cyan) and cell membrane (orange), derived from E-cadherin and phalloidin. *n* = 37 (E4.75), and 74 (E5.0) for *Itgb1*^+/+ or +/−^ and 16 (E4.75), and 26 (E5.0) for *Itgb1^−/−^* embryos examined from at least four independent embryo recovery experiments. (G-H) Angle measurement between the cell long axis and the normal vector to the tissue boundary, represented as violin plots with individual data points overlaid. Each plot shows the distribution of angles for EPI cells in contact with the ExE-boundary (G) and VE-boundary (H). Dot colors indicate the embryo stage at the collection. Data are grouped by EPI cell number, with group means shown by red bars. *n* = 1 (E4.5), 17 (E4.75), and 24 (E5.0) for *Itgb1*^+/+ or +/−^ and 2 (E4.5), 14 (E4.75), and 12 (E5.0) for *Itgb1^−/−^* embryos analyzed from at least four independent embryo recovery experiments Scale bars, 20 µm. See also Figure S3.

To experimentally test the functional role of boundary anchoring in EPI cell alignment, we genetically perturbed ECM deposition. At the VE-boundary, major laminin chains, laminin α1, β1, and γ1, were present (Figures S4A and S4C), though *Lama1* and *Lamb1* knockout embryos develop normally until E5.5, presumably due to functional compensation (Jeffrey H. Miner et al. 2004; J. H. Miner, Cunningham, and Sanes 1998). Genetic studies have demonstrated the essential role of laminin γ1 and integrin β1 in the egg-cylinder morphogenetic process (Smyth et al. 1999; Stephens et al. 1995), in agreement with recent studies using embryo models (Bedzhov and Zernicka-Goetz 2014; Molè et al. 2021). Here, we showed that EPI cells in *Lamc1*^−/−^ embryos were oriented tangentially to the VE-boundary, similar to their alignment at the ExE-boundary, in contrast to *Lamc1*^+/+^ or *Lamc1*^+/−^ controls (Figures 5B-5D). Moreover, the VE boundary in *Lamc1*^−/−^ embryos lacked active integrin β1 signal (Figure S3A), indicating that laminin γ1 is required for EPI cells to orient perpendicularly, through integrin β1-mediated anchoring. Tissue-scale analysis further revealed a loss of the polar alignment in *Lamc1*^−/−^ embryos, suggesting a disruption in EPI development (Figure 5E).

Furthermore, EPI cells in *Itgb1*^−/−^ embryos were oriented tangentially to the VE-boundary, unlike *Itgb1*^+/+^ or *Itgb1*^+/−^ littermates (Figures 5F-H), confirming the essential role of integrin β1 in guiding EPI cell orientation at the VE-boundary. Analysis of lumen nucleation by immunofluorescence of phospho-ERM showed that apical domain convergence was lost in *Itgb1*^−/−^ embryos (Figures S3B), consistent with the theoretical prediction for the lack of topological defects in the absence of the boundary anchoring that prefers perpendicular alignment (Figure 5A).

Together, these results demonstrate that integrin β1-laminin γ1-mediated adhesion is essential for EPI cells to form the polar alignment perpendicular to the VE-boundary, and for EPI tissue ordering and lumen formation.

### EPI tissue pattern facilitates activation of ERK pathway in EPI cells

The reduced EPI cell numbers observed in *Lamc1*^−/−^ and *Itgb1*^−/−^ embryos (see Figure 5) prompted us to test whether proper EPI cell alignment may be linked to cell differentiation and/or proliferation through activation of signaling pathways, such as the extracellular signal-regulated kinase (ERK) cascade. Immunofluorescence for phosphorylated ERK1/2 (pERK) in E4.75 – 5.5 embryos showed a progressive increase in the overall pERK signal within the EPI with notable cell-to-cell heterogeneity, in addition to those in ExE cells (Figures 6A and 6B) (Christodoulou et al. 2019; Kruger et al. 2024). Moreover, disruption of EPI cell alignment by 16-hours treatment with collagenase that selectively degrades collagen IV at the VE-boundary (Figure 6C) resulted in a reduction of the overall pERK levels in comparison to controls (Figures 6D and 6E). Together, these findings show that EPI tissue patterning promotes activation of ERK signaling in EPI cells, highlighting the critical role of cell alignment patterning in regulating signaling pathways crucial for cell differentiation and proliferation.

**Figure 6.**
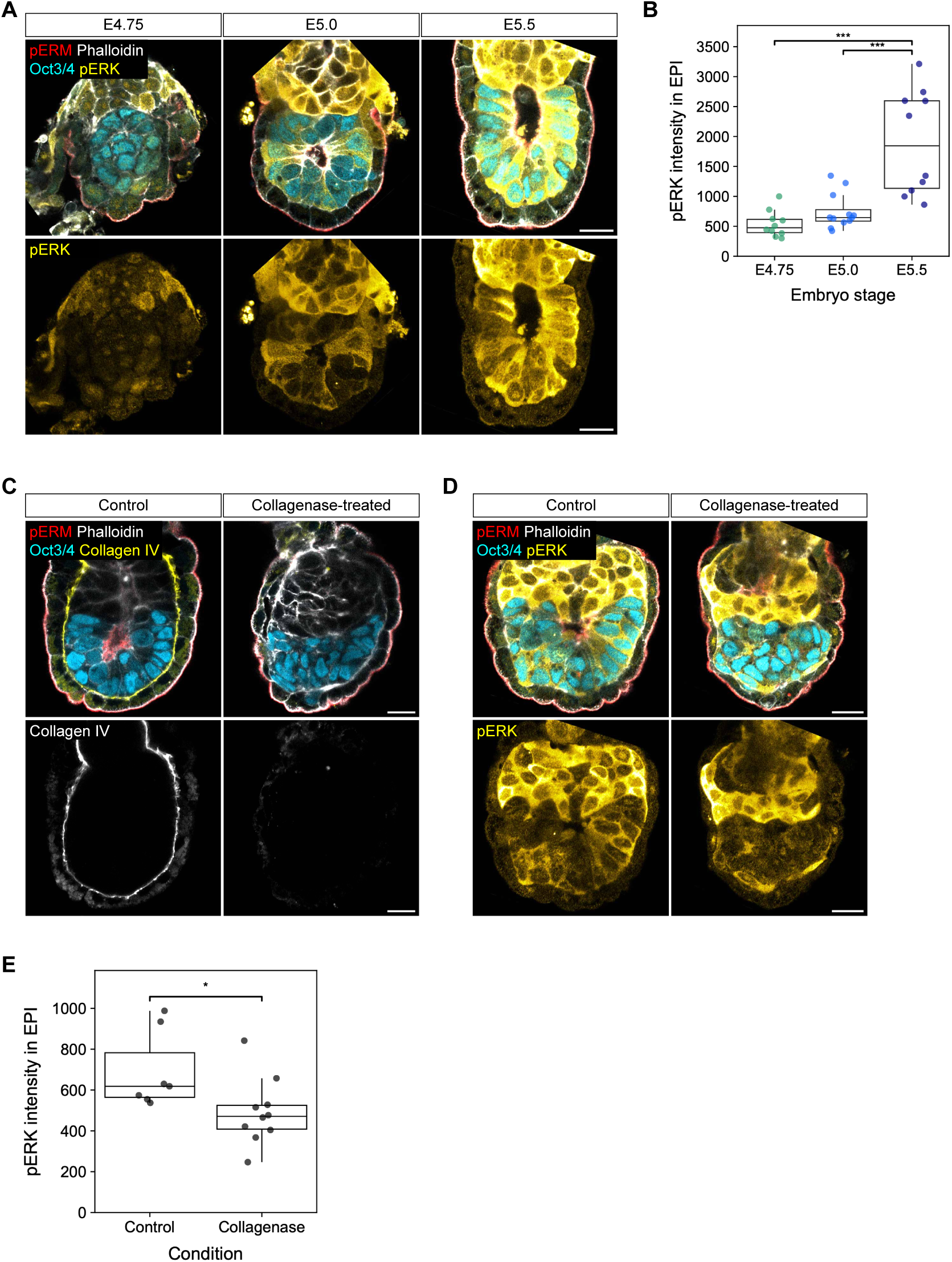
EPI tissue patterning facilitates ERK activation in EPI cells. (A) Immunofluorescence images of representative embryos from E4.75 to E5.5, stained for pERM (apical, red), phalloidin (white), Oct3/4 (EPI, cyan) and pERK (yellow). *n* = 21 (E4.75), 18 (E5.0), and 13 (E5.5) embryos analyzed from three independent embryo recovery experiments. (B) Quantification of mean pERK signal intensity within the EPI tissue. *n* = 10 (E4.75), 12 (E5.0), and 10 (E5.5) embryos measured from three independent embryo recovery experiments. (C) Immunofluorescence images of representative control and 500 μg/mL collagenase-treated embryos, stained for pERM (apical, red), phalloidin (white), Oct3/4 (EPI, cyan) and collagen IV (yellow). *n* = 8 (control), and 8 (collagenase-treated) embryos from two independent embryo culture experiments. (D) Immunofluorescence images of representative embryos cultured for 16 hours in control and 500 μg/mL collagenase-containing medium, stained for pERM (apical, red), phalloidin (white), Oct3/4 (EPI, cyan) and pERK (yellow). *n* = 14 (control), and 17 (collagenase-treated) embryos cultured from three independent experiments. (E) Quantification of mean pERK signal intensity within the EPI tissue. *n* = 7 (control) and 10 (collagenase-treated) embryos measured from three independent experiments.

## Discussion

This study reports that boundary-induced cell alignment drives the emergence of polarized architecture in the mouse EPI tissue. Our experiments, in combination with a theory of polar fluid ordering (Guruciaga et al. 2024), consistently show that cell-ECM interactions guide the EPI cell alignment, which in turn facilitates the functional maturation of the EPI to allow lumen formation and ERK activation.

The specific boundary condition defined by the localized expression of laminin at the VE-boundary is essential for EPI cell alignment. This finding is in agreement with earlier studies showing the role of integrin β1 and laminin γ1 in embryonic development through E5.5 (Smyth et al. 1999; Stephens et al. 1995; Sutherland, Calarco, and Damsky 1993). Moreover, we demonstrate that the differential expression of ECM components between the VE- and ExE-boundaries determines the characteristic EPI tissue architecture in the egg-cylinder, indicating the importance of the embryonic-extraembryonic tissue interface. Our theoretical framework, based on the physics of boundary-induced alignment, successfully captures the complex changes in cellular organization over time, despite its simplicity and reliance on just two key material parameters. Using 3D quantifications of cell elongation over developmental stages, we identified how the biophysical parameters associated with cell-boundaries and cell-cell interactions of the EPI tissue change over the course of development.

Our study couples changes in tissue boundary properties to the formation of lumen initiation sites within the EPI tissue (Guruciaga et al. 2024). These sites appear where the predicted orientation field features topological defects (Guruciaga et al. 2024) – localized singularities that determine the collective properties of ordered systems and guide diverse biological functions, including cell extrusion, protrusion, and morphogenesis (Guillamat et al. 2022; Kawaguchi, Kageyama, and Sano 2017; Saw et al. 2017). The heterogeneous boundary conditions of the EPI create lumen-inducing 3D topological defects when anchoring strength overcomes the effects of bulk interactions, suggesting that lumen formation coordinates the maturation of the tissue-ECM interface. Specifically, we show both *in silico* and *in vivo* that the ExE-boundary brings the nucleating lumen position closer to the ExE, thereby potentially contributing to embryo symmetry breaking through lumen-associated key signaling pathways, such as Nodal (Y. S. Kim et al. 2021) and BMP (Zhang et al. 2019).

While the role of FGF-ERK signaling has been studied in the EPI-PrE segregation in the blastocyst (Chazaud et al. 2006; Kang, Garg, and Hadjantonakis 2017; Molotkov et al. 2017; Ohnishi et al. 2014; Pokrass et al. 2020) and in the ExE in post-implantation development (Christodoulou et al. 2019; Kruger et al. 2024), its activity in the EPI in the peri-implantation embryo remained elusive. Our findings for the ERK signaling in the EPI are consistent with studies in ES cells that suggested the role of FGF-ERK signaling in driving the transition from naïve to formative pluripotency (Arekatla et al. 2023; Burdon et al. 1999; Kunath et al. 2007; Mulas et al. 2024). This suggests that the exit from the naïve state is coupled to EPI tissue organization during peri-implantation development (Shahbazi et al. 2017). Furthermore, cell-to-cell heterogeneity in ERK activation is in line with our previous observation of Dusp4 (Ichikawa et al. 2022), which negatively feeds back to FGF signaling (Azami et al. 2019). These findings provide a framework for future investigation into the regulatory mechanisms and dynamics of ERK activation, and exit from naïve pluripotency in the peri-implantation EPI.

During the EPI patterning in the peri-implantation mouse embryo, EPI cells undergo dynamic cellular changes, including elongation and apico-basal polarization (Bedzhov and Zernicka-Goetz 2014; Ichikawa et al. 2022; Y. S. Kim et al. 2021; Molè et al. 2021). Due to technical challenges in tracking polarity markers over time in the developing EPI, our current analysis infers the magnitude of polarity alignment from 3D cell shape measurements. Future studies will be valuable to gain insights into the contribution of cell polarization dynamics and its potential role in cell elongation during this tissue patterning process.

## Acknowledgments

We are grateful to the members of the Hiiragi group for valuable discussions and comments on the manuscript. Special thanks to Lidia Pérez and Miyuki Yoshida for their technical support. We thank Antoine Diez for assistance with mathematical analysis. We appreciate the animal care support provided by the Institute of Laboratory Animals, Graduate School of Medicine and the equipment support from the Single-Cell Genome Information Analysis Core (SignAC) at the Institute for the Advanced Study of Human Biology (ASHBi), Kyoto University. We thank Spyros Goulas for critical reading and constructive suggestions on the manuscript. We thank Hironobu Fujiwara for providing the Col4a2-eGFP mouse line. T.I. and S.P. are supported by Japan Society for the Promotion of Science (JSPS) KAKENHI (grant numbers JP25K18475 and JP24K16962, respectively). P.C.G. was supported by the EMBL Interdisciplinary Postdoctoral Fellowship (EIPOD4) programme under Marie Sklodowska-Curie Actions Cofund (grant agreement 847543) and an Add-on Fellowship for Interdisciplinary Life Science from the Joachim Herz Stiftung. S.H. was supported by the ASHBi Financial Support Program for International Graduate Students. The Hiiragi group is supported by the Graduate School of Medicine and ASHBi, Kyoto University, the Hubrecht Institute, Japan Society for the Promotion of Science (JSPS) KAKENHI (grant numbers JP21H05038 and JP22H05166), the European Research Council (ERC) Advanced Grants (“SelforganisingEmbryo” grant agreement 742732 and “COORDINATION” grant agreement 101055287), and Stichting LSH-TKI (LSHM21020). The Erzberger group is supported by the EMBL.

## Author contributions

Conceptualisation, T.I., P.C.G., A.E., T.H.;

Methodology, T.I., P.C.G., S.H., S.P., A.E., T.H.;

Software, P.C.G., S.P., A.S.;

Validation, T.I., P.C.G.;

Formal Analysis, T.I., P.C.G., S.H., S.P., M.M., M.H., A.S., S.Y.;

Investigation, T.I., S.H., M.M., M.H.;

Resources, T.I., A.E., T.H.;

Data Curation, T.I., S.H., S.P., A.S.;

Writing – Original Draft, T.I.;

Writing – Review & Editing, T.I., P.C.G., S.P., A.E., T.H.;

Visualisation, T.I., P.C.G., S.H., S.P.;

Supervision, T.I., A.E., T.H.;

Project Administration, T.I., T.H.;

Funding Acquisition, T.I., S.P., T.H.

## Additional information

Document S1. Figures S1-S3 and Table S1

Video S1. Light-sheet live-microscopy reveals EPI cell orientation dynamics, related to Figure 1. Time-lapse images of a Sox2-Cre;mT embryo developing in 3D-geec. Note that the equatorial plane of the embryo is shown in Figure 1E. Green, memb-GFP in EPI cells; magenta, memb-Tomato in other cell types. Time, hours:minutes, Scale bars, 20 µm.

## Methods

### Mouse work

All animal work was performed in the Institute of Laboratory Animals (ILA), with permission from the Animal Research Committee, Graduate School of Medicine (approved number, MedKyo 23065) and the Committee for Safety Control of Recombinant DNA Experiments, Kyoto University (approved number, 230029). ILA is operated according to the Regulations on Animal Experimentation at Kyoto University. All mice were maintained in specific pathogen-free conditions with a 14-10 hours light-dark cycle and used for experiments at the age of 8 to 30 weeks.

### Mouse lines and genotyping

The following mouse lines were used in this study: F1 hybrid strain between C57BL/6NCrSlc and C3H/HeSlc (B6C3F1/Slc) (Japan SLC) as wild-type (WT), mTmG (Muzumdar et al. 2007), *Sox2-Cre* (Hayashi et al. 2002), *Itgb1^tm1Efu (floxed)^* (Raghavan et al. 2000), *Lamc1^tm1Str (floxed)^* (Chen and Strickland 2003), and *Col4a2-eGFP* (Wuergezhen et al. 2025). To generate *Itgb1^+/−^* and *Lamc1^+/−^* mice, *Itgb1^floxed/floxed^*and *Lamc1^floxed/floxed^* females were crossed with *ZP3-Cre ^tg/+^* males, followed by crossing between *Itgb1^floxed/+^*;*Zp3-Cre^tg/+^* and *Lamc1^floxed/+^*;*Zp3-Cre^tg/+^*females and B6C3F1 males, respectively (E. J. Y. Kim, Sorokin, and Hiiragi 2022). Standard genotyping procedures were used to genotype transgenic mice (for primers and PCR product sizes, see Table S1).

### Embryo recovery

To obtain mouse embryos, mice were naturally mated, and the midpoint of the light period on the day when a vaginal plug was detected was defined as embryonic day 0.5 (E0.5). Recovery of embryos was performed under a stereomicroscope (Zeiss, Stemi 508) equipped with a thermo plate (Tokai Hit, TPi-STMX) heated at 37°C. Peri- and post-implantation embryos were recovered from dissected uteri in dissection medium (DMEM (Gibco, 11880028) supplemented with 15% heat-inactivated FBS (PAA, A15-080), 2 mM GlutaMAX (Gibco, 35050061), 10 mM HEPES (Sigma, H0887), 25 units/mL Penicillin and 25 µg/mL Streptomycin (Gibco, 15070063)). Peri-implantation embryos loosely adherent to the uterine luminal epithelium were isolated by opening along with the mesometrial side of the uterus, followed by a gentle touch using fine forceps (Dumont, No.5). Sites of embryo adherence can be identified by locally expanding tissue undergoing decidualization. Recovery of post-implantation embryos was performed as described previously (Behringer et al. 2013). The Reichert’s membrane of the post-implantation embryos was removed using Dentronics No.32 needles (Handaya, HS-2739B) for subsequent immunofluorescence staining. Recovered embryos were handled using an aspirator tube (Sigma, A5177) equipped with a pulled glass micropipette, 100 μL (Drummond, 2-000-1000), and cultured in an incubator (PHC, MCO-170AICUV) with a humidified atmosphere of 5% CO_2_ at 37 °C.

### 3D-gel embedded embryo culture (3D-geec)

3D-geec was performed as described previously (Ichikawa et al. 2022). Briefly, mural trophectoderm (mTE) was microsurgically removed from E4.5 embryos using Dentronics No.32 needles. mTE-removed embryos were then embedded in the gel droplet composed of 3.0 mg/mL growth factor reduced Matrigel (Corning, 356230, lot. 7345012) and 0.3 mg/mL Collagen I (Corning, 354236, lot.2055001), diluted in basal medium (advanced DMEM/F-12 (Gibco, 12634010) supplemented with 2 mM GlutaMAX, 25 units/mL Penicillin and 25 µg/mL Streptomycin). After solidification of the gel upon 30 minutes of incubation in the incubator, 50 µL pre-warmed IVC1 medium (Bedzhov and Zernicka-Goetz 2014) was added to cover the gel. To degrade collagen IV, NP-collagenase (Nippi, 892461) was added to the IVC1 medium at a concentration of 500 µg/mL.

### Single embryo genotyping

Transgenic mutant embryos were genotyped retrospectively after imaging. Single embryos were transferred using a mouth-pipette from the imaging dish into PCR tubes containing 10 μL lysis buffer composed of *Ex Taq* buffer (Takara, RR006A) supplemented with 0.2 mg/mL Proteinase K (Sigma, P2308). Embryos in the lysis buffer were incubated at 55°C for 1 hour and then at 96°C for 10 minutes. 5 μL of the genomic DNA lysate was subjected to PCR using *Ex Taq*.

### Immunofluorescence staining and imaging

Embryos were fixed with 4% paraformaldehyde (Fujifilm Wako, 166-23251) in PBS for 20 (E4.5-5.0) or 30 (E5.25-5.5) minutes at room temperature and subsequently permeabilized with 0.5% Triton X-100 (Nacalai, 12967-32) in PBS for 30 minutes at room temperature with gentle agitation. Embryos were incubated in blocking buffer (3% BSA (Sigma, A9647), 0.05% Triton X-100 in PBS) overnight at 4^°^C with gentle agitation. Embryos were then incubated with primary antibodies diluted in the blocking buffer overnight at 4^°^C or 2 hours at room temperature. After washing with the blocking buffer, embryos were further incubated with secondary antibodies diluted in the blocking buffer for 2 hours at room temperature. Phalloidin staining was simultaneously performed with the secondary antibody staining, using Alexa Fluor Plus 405 Phalloidin (Invitrogen, A30104) diluted at 1:400. Finally, stained embryos were transferred into PBS droplets overlaid with mineral oil, on a 35 mm glass base dish (IWAKI, 3970-035) for imaging.

Primary antibodies against Oct3/4 (Santa Cruz Biotechnology, sc-5279 AF647), E-cadherin (BD Biosciences, 560064), active integrin β1 (9EG7, BD Biosciences, 553715), and phosphorylated-ERK1/2 (p44/42 MAPK) (Cell Signaling, 4370) were diluted at 1:100. Primary antibodies against laminin (Novus-Biologicals, NB300-144), phosphorylated-ERM (Cell Signaling, 3726), and podocalyxin (R&D Systems, MAB1556) were diluted at 1:200. Laminin chain-specific antibodies were used as described previously (E. J. Y. Kim, Sorokin, and Hiiragi 2022). Secondary antibodies, donkey anti-rabbit IgG Alexa Fluor Plus 488 (Invitrogen, A32790), donkey anti-rabbit IgG Alexa Fluor Plus 555 (Invitrogen, A32794), and donkey anti-rat IgG Alexa Fluor Plus 488 (Invitrogen, A48269) were used at 1:200.

Images of immunostained embryos were acquired by an LSM980 microscope equipped with a C-Apochromat 40x/1.2 NA water immersion objective (Zeiss), with Airyscan 2 Multiplex CO-8Y mode. Raw Airyscan images were post-processed by ZEN blue software (Zeiss). The image voxel size after the Airyscan processing was 0.0823 x 0.0823 x 0.1600 µm^3^ (X, Y, Z).

### Light-sheet Live-imaging

3D-geec embryos were live-imaged using an inverted light-sheet microscope (Bruker, Luxendo, InVi SPIM), as described previously (Ichikawa et al. 2022). Briefly, embryos were embedded in a 15 µL gel mix within the V-shaped sample holder attached with transparent FEP foil, carefully positioned so that they are in proximity but do not attach to the foil that would disrupt morphogenesis via adhesion. After gel solidification, embryos were immersed in 75 µL IVC1 medium and further covered with 200 µL mineral oil to prevent evaporation. The sample holder was mounted in an environmentally controlled incubation box with 5% CO_2_ and 5% O_2_ at 37 °C.

InVi SPIM was equipped with a Nikon 25x/1.1NA water immersion detective objective and a Nikon 10x/0.3 NA water immersion illumination objective. The illumination plane and focal plane were aligned before the imaging session and maintained during the imaging. Images were taken every 20 min by a CMOS camera (Hamamatsu, ORCA Flash4.0 V2) with line-scan mode in LuxControl (Luxendo). The imaged volume was 212.99×212.99×200 µm^3^ with a physical voxel size of 0.104 x 0.104 x 1.000 µm^3^. The lasers and filters used were 488 nm and BP525/50, and 561 nm and LP561 to image GFP and tdTomato fluorophores, respectively. Exposure time for each plane was set to 30 ms.

### Simulations of surface-induced order in a polar fluid

We consider a polar fluid in a space Ω with volume *V*_0_ confined by the surface ∂Ω. This system can be characterized by a local order parameter **p**(**r**), which defines the global degree of order 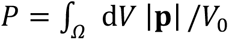 and minimizes the free energy functional

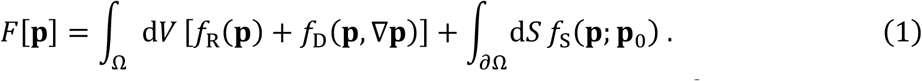

The first term corresponds to the bulk energy, with a restoring term *f*_R_ = *a*|**p**|^2^/2 with a ≳ 0 (i.e., the system is in the isotropic phase) and a distortion energy density 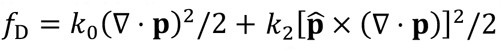 where 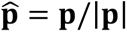 and *k*_0_, *k*_2_ penalize the splay and bend distortions (the twist contribution vanishes due to symmetry (Guruciaga et al. 2024)). The second term represents the surface energy, given by a weak anchoring interaction with *f*_S_ = *w*(**p** − **p**_0_)^2^/2 where **p**_0_ is the preferred value for the order parameter at the boundary. We consider cases where the confining surface is formed by two distinct surfaces, *S*_*⍺*_ (ExE-boundary) and *S*_*β*_ (VE-boundary), and we take 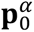 to be tangent to *S*_*⍺*_ and 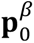 to be normal to *S*_*β*_. If the spatial coordinates are normalized by the characteristic length *R*_0_ = (3*V*_0_/4*π*)^1/3^, equation (1) can be rewritten in the rescaled space Ω′ as

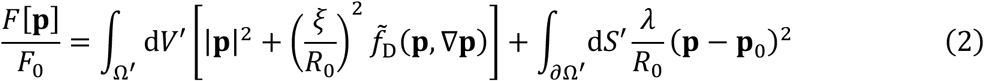

where 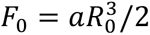 and 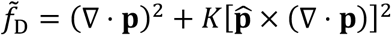 with *K* ≡ *k*_2_/*k*_0_. Parameters 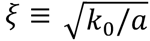 and *λ* ≡ *w*/*a* are, respectively, the correlation and anchoring lengths of the system, and tune the importance of the distortion and anchoring contributions.

In order to minimize the free energy (2), we implemented the finite-element method using the FEniCSx library DOLFINx (Scroggs et al. 2022) in Python3. Given the axial symmetry of the system, the mesh (with resolution 0.05) was defined in spherical coordinates in terms of *r* and *ϕ* only, corresponding to a constant-*ϕ* slice of the three-dimensional system. We computed the variation of equation (2) with respect to **p** in the direction of a test function *ψ* to derive its weak formulation. The resulting nonlinear problem was solved using a Newton solver with a relative tolerance of 10^−6^.

### Fitting geometrical parameters to embryo shape

We consider axis-symmetric confining surfaces like the one in Fig. 3B of the main text, formed by two spherical shells *S*_*μ*_, *μ* = *⍺*, *β* representing the ExE-EPI and VE-EPI interfaces, respectively. Each shell is centered at the point (0, *c*_*μ*_) on the symmetry axis and has a radius *R*_*μ*_ ≡ 1/***κ***_*μ*_, where ***κ***_*μ*_ is the curvature. They can be parameterized in spherical coordinates as

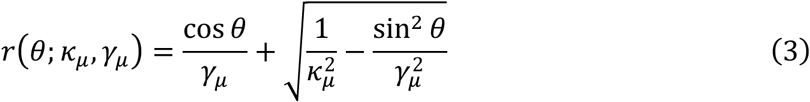

where *γ*_*μ*_ ≡ 1/*c*_*μ*_, with *θ* ∈ [0, *π*/2] for *μ* = *⍺* and *θ* ∈ [*π*/2, *π*] for *μ* = *β*, and *ϕ* ∈ [0,2*π*). In order to determine the best set of geometrical parameters *R*_*μ*_, *c*_*μ*_ for a given EPI shape, we fit equation (3) to ExE (*μ* = *α*) and VE (*μ* = *β*) boundary data as described in (Guruciaga et al. 2024). Briefly, we convert the *N*_*μ*_ experimental data points to spherical coordinates, 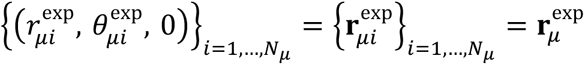, and minimize the cost function

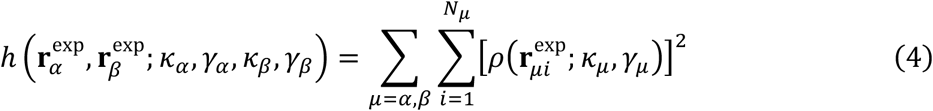

while enforcing the constraint *r*(*π*/2; ***κ***_*⍺*_, *γ*_*⍺*_) = *r*(*π*/2; ***κ***_*β*_, *γ*_*β*_). In equation (4),

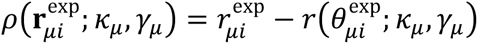 is the residual between the radius of experimental point *i* of boundary *μ* and the fitting function (3) evaluated at that point.

### Image analysis

Pre-processing for machine-learning-based membrane segmentation and signal intensity measurement were performed with Fiji (Schindelin et al. 2012). Membrane segmentation and custom model training were performed using Cellpose (Stringer et al. 2021; Pachitariu and Stringer 2022) GUI or CLI using a bash script. Manual correction of segmentation and cell tracking were performed with Napari (Sofroniew et al. 2024). Measurements of the angle between long axes of neighboring cells, measurements of the cell orientation, analysis of the correlation between the signal intensity and the cell orientation, and the tissue-scale alignment analysis were performed using custom Python scripts.

### Machine-learning-based membrane segmentation

The segmentation pipeline used to process the 3D images of the membrane signal consists of four steps. In the first step, the source 3D images were pre-processed to generate the membrane channel with isotropic voxel size. In the second step, custom segmentation models were developed on the Cellpose platform. In the third step, segmentation tasks were performed in batch mode using the Cellpose CLI. In the fourth step, the EPI membrane segmentation was manually selected and corrected in Napari. The details of the individual steps are described below.

#### 1. Pre-processing

Airyscan images were binned to 0.1647 x 0.1647 x 0.1600 µm^3^ by averaging 2 x 2 x 1 voxels, followed by isotropic transformations of voxels to a cube with a length of 0.1632 ± 0.0002 µm. To quantify spatial parameters, we set the length of the voxel to 0.1632 µm and ignored the associated error. Images acquired by light-sheet microscope were first cropped to remove background voxels as much as possible and binned to 0.208 x 0.208 x 1.000 µm^3^ by averaging 2 x 2 x 1 voxels, followed by isotropic scaling to 0.351 ± 0.001 µm.

Two channels for E-cadherin and phalloidin signals of the isotropically scaled Airyscan images were combined to generate a ubiquitous membrane channel by summing their signal intensities (Figures 1A and S1A). Similarly, two channels for mG and mT signals of the isotropically scaled light-sheet microscopy images were also used to generate a membrane channel (Figure 1E). These membrane channels were used as inputs for segmentation.

#### 2. Developing custom segmentation models

Since the pre-trained models provided by Cellpose 2.0 required an insuperable amount of manual correction for our 3D images, we developed custom segmentation models tailored for Airyscan images and light-sheet microscopy images independently.

To develop models for Airyscan images, pre-processed images and their corresponding ground truth masks were prepared. In brief, a pre-processed 3D image of an E5.0 stage embryo was resliced into three orthogonal planes (XY, YZ, and XZ), and five slices were extracted in each plane at a 50-slice interval across the entire image. The resulting slices were subjected to 2D segmentation using the Cellpose pre-trained model, CPx, followed by manual corrections of all cells in the slice using Napari. Following the instructions, two embryo datasets, each consisting of 15 pairs of slices and ground truth masks, were used to train the neural network, generating “AS_model_1” (Figure S1B). To further improve AS_model_1, we conducted additional training by incorporating two extra embryo datasets into the neural network, resulting in “AS_model_2”. While both models showed comparable performance on images that have high signal-to-noise ratios, AS_model_2 is more robust to the images with various signal-to-noise ratios. Therefore, we used AS_model_2 in this study (Figures 1 and 4).

Models for light-sheet microscopy images were developed using Cellpose with sparse annotations (Sugawara 2023). The combined membrane stacks at three different time points of a time-lapse images were subjected to 3D segmentation using the CPx model, followed by manual corrections of EPI cells using Napari. These ground-truth images were used to train the neural network iteratively, generating “LS_model_4”.

#### 3. Batch segmentation

For 3D segmentation of Airyscan images, segmentation parameters, such as flow_threshold, cellprob_threshold, and stitch_threshold, were set to default values, except for the cell diameter, which was set to 60 pixels. The segmentation model “AS_model_2” was chosen for this process.

For 3D segmentation of light-sheet microscopy images, segmentation parameters, such as flow_threshold, cellprob_threshold, and stitch_threshold, were set to default values, except for the cell diameter, which was set to 30 pixels. The segmentation model “LS_model_4” was selected for this process.

#### 4. Manual correction

Segmentation errors were manually corrected by referring to the original images, and EPI cells were selected and counted in Napari using a custom plugin (napari-labelselector, unpublished). The corrected segmentation masks were saved as ‘.tif’ files for further analysis.

### Neighboring cell angle measurement

Neighboring cell angle measurements were performed using Python 3.9 based on manually corrected voxel-based segmentation masks. The principal inertia vectors of a label were computed, and the angle between the long axes (primary components) of adjacent labels was measured as the neighboring cell angle. Neighboring cells were identified by dilating the label after binarization and computing the overlapping labels by element-wise multiplication. Angle data from the neighboring cell pairs with overlapping volumes smaller than 10 μm^3^ were excluded from the analysis. The frequency of angle distribution was normalized within each group based on EPI cell number.

### Mesh-based computations of polarity vectors, surface normals, and alignment

To analyze cell alignment and polarity within the EPI tissue, we first converted the corrected EPI segmentation into a mesh representation using the Python package *scikit-image* (van der Walt et al. 2014). Each segmented cell was reconstructed as an individual mesh, and a global mesh for the entire EPI region was generated. To ensure consistency in the analysis, all meshes were post-processed to correct face orientations, and only the largest connected component was retained. The final EPI mesh stored cell labels as vertex attributes, enabling cell-specific calculations. The boundary normal vector for a given cell was computed as the average normal vector of all mesh vertices from the EPI mesh associated with that cell. We then used the *trimesh* library (https://github.com/mikedh/trimesh) to compute various geometric and topological properties, including cell volume, surface area, centroid, and principal inertia vectors. Additionally, the centroid of the entire EPI region was determined to facilitate further spatial analyses.

The polarity vector was defined by the largest principal inertia component. Polarity vectors were flipped if the corresponding cell boundary normal was in the opposite direction or, for interior cells, if the vector was pointing towards the EPI center. The scaled polarity vector was the polarity vector multiplied by s = 1 − λ_1_/(λ_2_ + λ_3_), where λ_1_ ≥ λ_2_ ≥ λ_3_ were the sorted principal inertia components. Cell orientation to the tissue boundary was computed for all cells touching the boundary as the angle between the polarity vector and the normal vector ranging from 0° to 90°, where 0° indicates the perpendicular orientation while 90° the parallel.

### Determination of distal-proximal axis

Imaging data was annotated with at least six points in Napari, through which a regression plane approximately separates the boundary between the ExE and VE regions. One point was annotated to denote the distal tip of the EPI. We defined a rotation axis as a line that passes through the EPI centroid and the annotated tip point. Two points were also annotated to indicate the transverse edge of the EPI, the mean position of which was used to establish a rotation plane at angle 0°.

### Weighted nematic average over rotational slices

To construct a 3D nematic vector field representing average cell alignment, we first reoriented the coordinate system such that the *z*-axis aligned with the distal-proximal axis, previously annotated. The spatial coordinates were then rescaled to normalize the EPI volume to unity. At each spatial location **r** = (*x*, *y*, *z*), we extracted the major principal inertia vector *v*(**r**) and its corresponding inertia components *λ*_1_ ≤ *λ*_2_ ≤ *λ*_3_ from the occupying cell mesh. The elongation axis was then defined as *v*′(**r**) = *η*(**r**)*v*(**r**) where the shape anisotropy factor 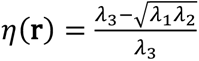 ranged from 0 for spheres up to 1 for infinitely elongated shapes. Next, we defined M = 36 equidistant angles *θ*_*m*_, generating rotational slices around the *z*-axis. For each slice, we used the rotation matrix *R*(*θ*_*m*_) which maps from the rotational slice to the *xz*-plane to obtain elongation axes within the *xz*-plane as 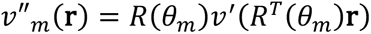. We then computed the weighted Landau-de Gennes Q-tensor, defined as 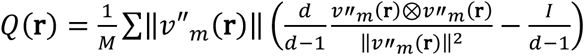 for *d* = 3.Finally, we took its principal eigenvector *V*(r) as the (unsigned) director field and the corresponding eigenvalue *Λ*(**r**) as the strength of the nematic alignment. This yields the average cell alignment vector *w*(**r**) = *Λ*(**r**)*V*(**r**). For visualization in 2D, we evaluated this quantity over a uniform grid in the xz-plane, displaying only the in-plane components (*w*_*x*_, *w*_*z*_) while additionally using the strength of the nematic alignment *Λ*(**r**) as a heatmap.

### Generation of the rotationally averaged embryo boundary curve

To obtain a rotationally averaged embryo boundary, we first computed the intersection of the EPI mesh with the M rotational slice planes defined by angles *θ*_*m*_. These boundary curves were converted into polar coordinates, with the EPI centroid as the origin. The curves were resampled at equidistant angles, and their radial distances were averaged across all slices to produce a smooth, rotationally averaged boundary. Finally, the averaged boundary was transformed back into Cartesian coordinates for visualization in the xz-plane.

### Average nematic magnitude

To quantify the overall nematic alignment of one embryo, we defined the average nematic magnitude as 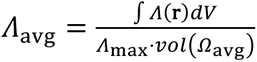 where *Ω_avg_* is the 3D volume enclosed by the averaged embryo boundary and *Λ*_max_ denotes the maximal value of the nematic strength factor over all embryos and all positions.

### Cross-embryo averaged nematic averages and boundary curves

For the cross-embryo nematic alignment field, the Q-tensors were averaged across all embryos as 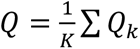 where *K* is the number of embryos and *Q*_*k*_ denotes the previously computed Q-tensors of individual embryos. Similarly, we compute the averaged embryo boundary by averaging with respect to all rotational slices from multiple embryos.

### Fitting the material length scales to the experimental cell elongation field

The nematic alignment tensor captures the cell’s orientation coherence, but not the polarity direction; its magnitude, however, can provide an unsigned estimation of the polarity magnitude, assuming that tail-head inversion is negligibly rare. We define the experimental polarity magnitude as *p*_exp_ ≡ *Λ*/*Λ*_max_, where the strength of the nematic alignment *Λ* is normalized by its maximum across stages *Λ*_max_. This quantity allows us to define the experimental global order as 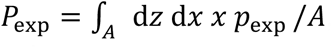, where *A* is the area defined by the average EPI boundaries and the *z*-axis.

In order to determine the model parameters that best characterize a given embryo stage, we fit the average EPI shape as described before, and minimize the free energy functional (2) in such system for the shown range of mechanical parameters *ξ*/*R*_0_ and *λ*/*R*_0_, keeping *K* = 10^−2^ constant (Guruciaga et al. 2024) to obtain the corresponding theoretical field with magnitude *p*. In order to compare the experimental and theoretical results, we define a set of *N* points **r**_*i*_ belonging to the intersection of the theoretical and experimental embryo shapes. These points must be at a distance greater than 0.05 from the symmetry axis and greater than *d* from the closest boundary, were *d* is given by the mean standard deviation of the experimentally determined average boundary. In this way, we exclude regions where the average experimental field has low statistics due to the different shapes of the individual embryos. Finally, we calculate the cost function

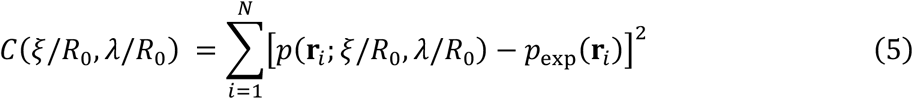

and identify the set of relative correlation and anchoring lengths, *ξ*/*R*_0_ and *λ*/*R*_0_, where it attains its minimum value.

### Statistical analysis and data reproducibility

Data were analyzed using R [Version 2023.09.1+494] or Python v3.9. Data normality was assessed using Shapiro-Wilk test. The EPI cell number distribution followed the normal distribution while the cellular morphological parameter data did not. Non-parametric tests were employed for data that did not meet the assumptions of normality. The notation for different p-values is as follows: ns, p > 0.05; *, p < 0.05; **, p < 0.01; ***, p < 0.001; ****, p < 0.0001.

Comparisons of the EPI cell number between various embryo stage groups were conducted using one-way ANOVA followed by Tukey’s post hoc test. Comparisons of all the cellular morphological parameters between groups were done using Wilcoxon-Mann-Whitney test (for comparing two groups) or Kruskal-Wallis test (for comparing three more groups).

### Materials Availability

All unique/stable reagents generated in this study are available from the corresponding authors with a completed Materials Transfer Agreement.

### Data and Code Availability

All datasets/codes generated during this study are available upon request.

