## Supplementary information for "Boundary-guided cell alignment drives mouse epiblast maturation"

### Figure S1

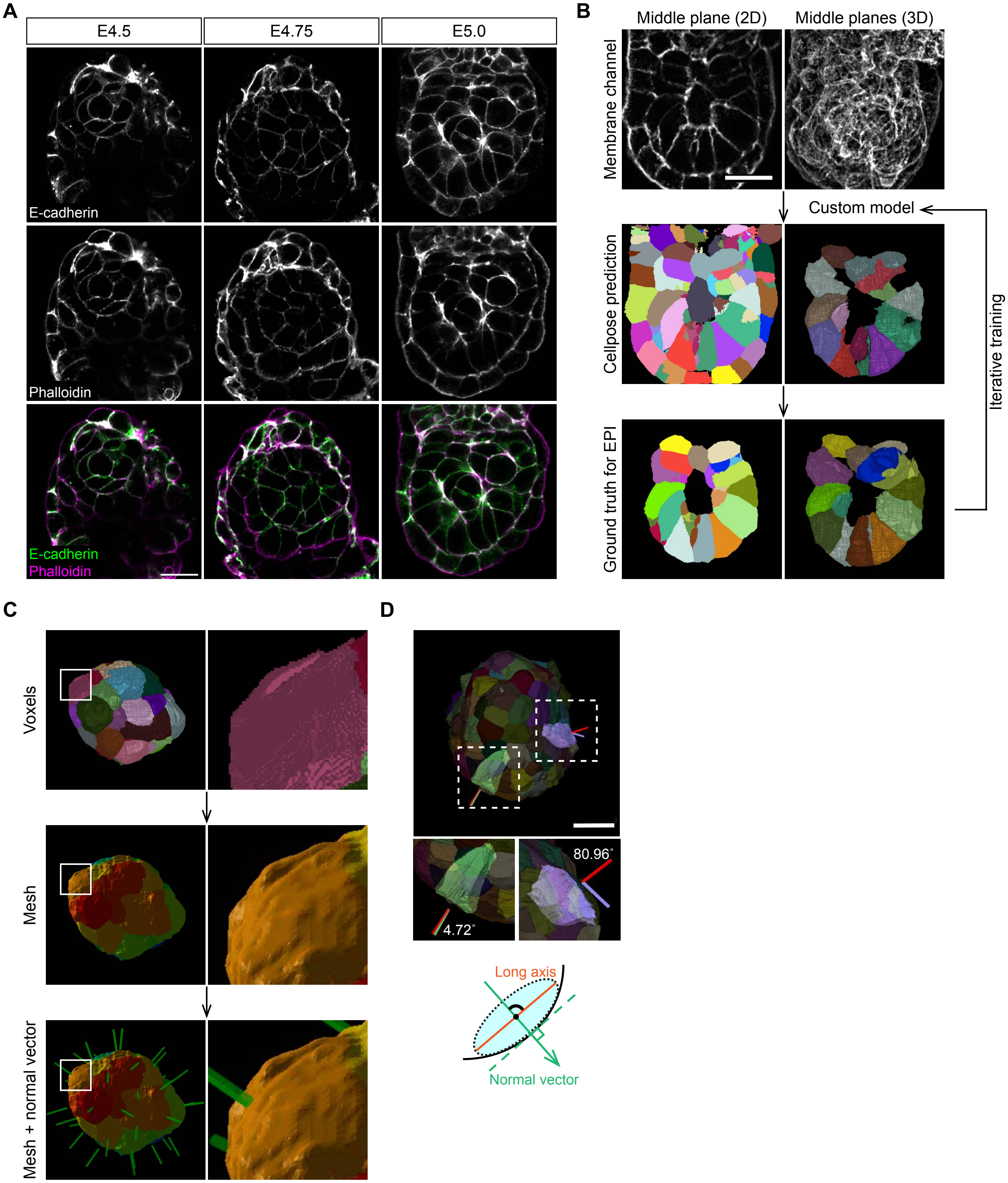


**Figure S1. Systematic analysis of EPI cell morphological parameters using 3D membrane segmentation.**

(A) Immunofluorescence images of representative embryos from E4.5 to E5.0, stained for E-cadherin and phalloidin (actin) to generate cell membrane signals.

(B) Workflow of 3D membrane segmentation using Cellpose. Membrane channel images generated in (A) (top row) were processed with Cellpose 2.0 (middle row). Manual corrections of all EPI cells using Napari were used for custom model training.

(C) Surface mesh conversion from voxel-based segmentation images. Each cell was reconstructed as an individual mesh, while the entire EPI tissue was converted to a global mesh. For cell orientation measurement, the average normal vector of all corresponding mesh vertices was computed (green bars) (see Methods).

(D) EPI cell orientation to the tissue boundary was measured as an angle between the normal vector calculated in (C) and the major axis of the cell (see Methods).

Scale bars, 20 μm.

See also Figure 1.

**Figure S2**

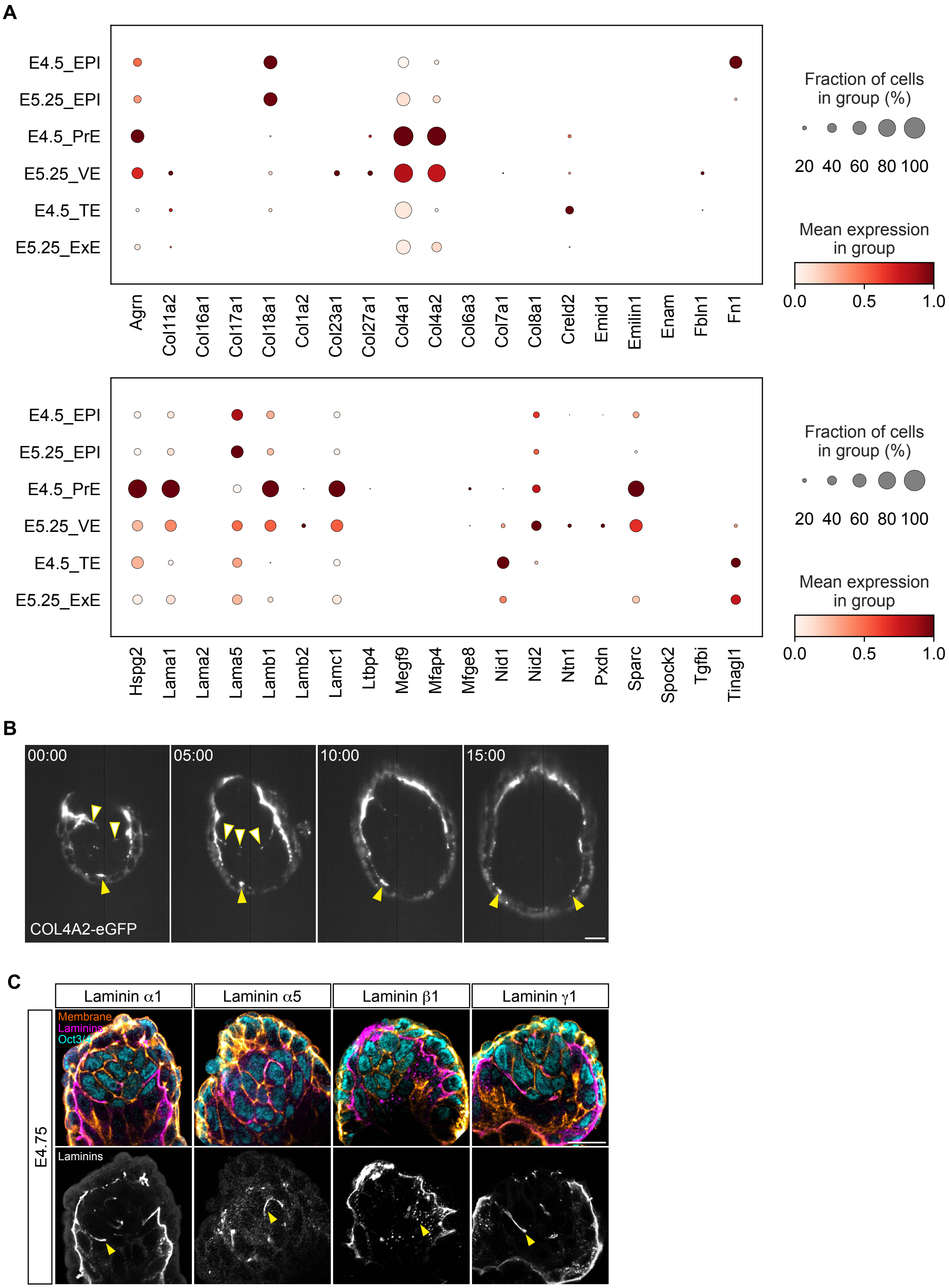
A
E4.5_EPI
E5.25_EPI
E4.5_PrE
E5.25_VE
E4.5_TE
E5.25_ExE
Agrn
Col11a2
Col16a1
Col17a1
Col18a1
Col1a2
Col23a1
Col27a1
Col4a1
Col4a2
Col6a3
Col7a1
Col8a1
Creld2
Emid1
Emilin1
Enam
Fbln1
Fn1
Fraction of cells in group (%)
20 40 60 80 100
Mean expression in group
0.0 0.5 1.0
Hspg2
Lama1
Lama2
Lama5
Lamb1
Lamb2
Lamc1
Ltbp4
Megf9
Mfap4
Mfge8
Nid1
Nid2
Ntn1
Pxdn
Sparc
Spock2
Tgfbi
Tinagl1
B
00:00
05:00
10:00
15:00
COL4A2-eGFP
C
Laminin α1
Laminin α5
Laminin β1
Laminin γ1
E4.75
Membrane
Laminins
Oct3/4
Laminins

**Figure S2. Characterisation of ECM molecules at the tissue boundary.**

(A) Single-cell transcriptome data analysis using Bondarenko et al. 2023. Gene ontology terms for ECM molecules were used to search expression levels in single-cells isolated from E4.5 and E5.25 embryos. EPI, PrE, VE, TE and ExE were annotated by Leiden clustering and known markers.

(B) Time-lapse images of representative Col4a2-eGFP embryos developed in 3D-geec. Time is shown as hh:mm, with t=00:00 marking the start of imaging. Signals accumulated at the VE-boundary (yellow arrowhead) while diminished at the ExE-boundary (white arrowhead). $n$ = 3 embryos.

(C) Immunofluorescence images of representative E4.75 embryos stained for laminin $\alpha$1, laminin $\alpha$5, laminin $\beta$1, and laminin $\gamma$1, together with cell membrane (E-cadherin and phalloidin), and Oct3/4 (EPI).

Scale bars, 20 μm.

See also Figure 4.

### Figure S3

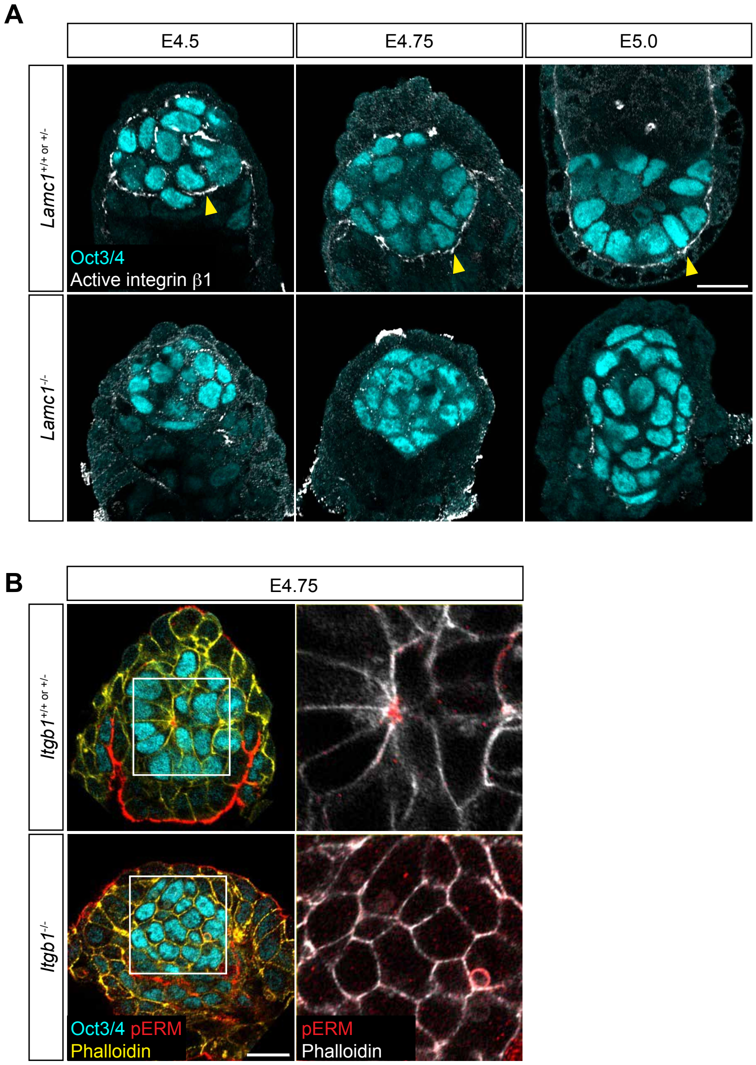


**Figure S3. Impacts of *Lamc1* and *Itgb1* genetic perturbations on apico-basal polarisation.**

(A) Immunofluorescence images of representative $Lamc1^{+/+ \text{ or } +/-}$ and $Lamc1^{-/-}$ embryos from E4.5 to E5.0, stained for Oct3/4 (EPI, cyan) and active integrin β1 (grey). Yellow arrowheads indicate signals at the VE-boundary.

(B) Immunofluorescence images of representative $Itgb1^{+/+ \text{ or } +/-}$ and $Itgb1^{-/-}$ embryos at E4.75, stained for Oct3/4 (EPI, cyan), phospho-ERM (red), and phalloidin (yellow).

Scale bars, 20 µm.

See also Figure 5.

| **Mouse Line** | **Primer ID** | **Primer Sequence** | **PCR Product Size, bp** |
|---|---|---|---|
| mTmG | oIMR7318 | CTCTGCTGCCTCCTGGCTTCT | WT allele, 330; Knock-in allele, 250 |
| | oIMR7319 | CGAGGCGGATCACAAGCAATA | |
| | oIMR7320 | TCAATGGGCGGGGGTCGTT | |
| Sox2-Cre | Sox2WT_Fw_37581 | CTTGTGTAGAGTGATGGCTTGA | WT allele 207 bp; Tg allele, 165 |
| | Sox2WT_Rv_37582 | TAGTGCCCCATTTTTGAAGG | |
| | Sox2Cre_Fw_37583 | CCAGTGCAGTGAAGCAAATC | |
| *Itgb1*(del) | Itgb1 del_F | TGAATATGGGCTTGGCAGTTA | WT allele, 900; Knock-out allele, 400 |
| | Itgb1 del_R | CCACAACTTTCCCAGTTAGCTCTC | |
| | Itgb1 del floxed | CGCAGAACAATAGGTGCTGAAATTAC | |
| *Lamc1*(del) | Lamc1_1 | AAAGAAGCAGAGTGTGGGGG | WT allele, 408; Knock-out allele, 700 |
| | Lamc1_2 | TGGCCTTTTCAACCCTGGAA | |
| | Lamc1_3 | GCCTTCTATCGCCTTCTTGAC | |

**Table S1. Genotyping primers and PCR product sizes.**
